# Successful gall induction by *Diplolepis rosae* and *D. mayri* on the sweet briar (*Rosa rubiginosa*) under laboratory conditions

**DOI:** 10.1101/2023.11.22.568036

**Authors:** Zoltán László, Mátyás Biró, Constantin-Teodor Iordache, Borbála Macalik, Marco Nicula, Bálint Szilágyi, Dorina Podar

## Abstract

1. Plant galls are unique outgrowths caused by various organisms, including insects, serving as nourishment for the inducer’s larvae. Despite the taxonomists and ecologists attempts to elucidate the mechanisms behind plant gall formation, its understanding is still incomplete. Modern genetic techniques allow in depth analysis of the molecular processes, but variations across species entangle the analysis. Establishing laboratory-friendly plant-gall inducer communities is crucial, yet past attempts have faced challenges.
2. Our study aimed to create a sustainable laboratory community involving wild roses (*Rosa* sp.) and as gall-inducing insects rose gall wasps belonging to the genus *Diplolepis*. Controlled indoor conditions were optimized for plant growth. Wild roses were transplanted, then exposed to gall inducers, and monitored.
3. Successfully initialized gall growth was measures and analysed, revealing insights into the impact of plant vigour on gall size.
4. Our study successfully established a novel laboratory community for further research on gall formation mechanisms.

## Introduction

Plant galls are unique structures caused by various organisms like bacteria, fungi, nematodes, and arthropods. Gall inducers force plants to develop novel tissues and organs, serving as nourishment for their offspring (Stone & Schönrogge, 2003).Taxonomists and ecologists have long been captivated by plant galls, even with incomplete comprehension of their origin. Deciphering the induction mechanism has been the primary goal for plant gall scholars since their discovery. Contemporary genetic and molecular techniques enable a profound exploration of the molecular processes driving gall formation (Giron *et al*., 2016). However, unravelling these mechanisms will be a prolonged endeavour due to the diverse evolutionary occurrences of gall formation in the animal kingdom, implying distinct mechanisms across taxa (Gätjens-Boniche, 2019). Consequently, identifying suitable plant and gall inducer model communities remains crucial.

Selecting a suitable model community of plant-gall inducers is decisive for understanding gall formation, requiring in-depth insights at the molecular level. To achieve this, -omics techniques (such as genomics, transcriptomics, proteomics) or their combinations prove invaluable. These methods involve rapidly cooling freshly collected samples to -80°C to preserve transient substances within tissues and cells. Achieving rapid cooling within hours or days depends on the distance of sampling. However, using liquid nitrogen in the field for on-site cooling to -80°C is both expensive and potentially hazardous. To prevent degradation and maintain cellular components, a practical approach is to establish the model community near a laboratory refrigerator. This approach also offers other benefits: a precise understanding of gall formation’s developmental stages, enabling precise monitoring and sampling; and controlled environmental conditions, enhancing omics analyses’ accuracy by reducing the impact of external variations.

Within arthropods, certain gall-inducing insects exhibit specialised connections to specific host plant species or groups, such as oak and rose gall-forming cynipids (Hymenoptera: Cynipidae). Limited efforts have been made in cultivating gall inducer communities and their host plants in laboratory settings. These attempts had limited success due to the need for continuous presence of gall inducer larvae throughout the developmental cycle to sustain plant tissue growth (Rohfritsch, 1971), or because they focused on partial aspects, excluding the complete community (Barrett *et al*., 1998). While a laboratory-friendly model galling community involving the Hessian fly (*Mayetiola destructor*) and wheat (*Triticum* spp.) exists, the induced galls lack distinct structural features (Stuart *et al*., 2012), leaving gaps in understanding gall-forming mechanisms from this case. Recently, a lab-maintainable community was established, featuring a weevil (*Smicronyx madaranus*) inducing galls on field dodder (*Cuscuta campestris*), a parasitic plant on various hosts (Murakami *et al*., 2021). However, this community involves herbaceous plants and insects, therefore, the exploration of woody plants like shrubs and trees within this context is still lacking.

Our study aimed to cultivate *Diplolepis rosae* (Linnaeus, 1758) and *D. mayri* (Schlechtendal, 1877) galls under laboratory conditions by transferring young wild roses from the field and subsequently infecting them with gall inducers. We present a novel laboratory community for breeding and sustaining gall-forming insects, paving the way for in-depth plant physiological and molecular analyses of diverse gall-inducing mechanisms. This community comprises the woody shrub sweet briar (*Rosa rubiginosa* Linnaeus, 1771) as the host plant and the well-known mossy rose gall (*D. rosae*) along with its sister species *D. mayri*.

## Material and Methods

To emulate the natural habitat of wild roses, we designed an indoor environment optimized for plant growth in the laboratory. Controlled temperature (20°C) and humidity upheld optimal plant conditions. The 12-hour day/night photoperiod mirrored natural light cycles, essential for circadian-influenced plant growth. LED FITO Full Spectrum and neutral/cold white lamps covered a range of wavelengths, accommodating various developmental needs. This integrated approach, combining air conditioning, humidity control, lighting precision, and LED technology, fostered controlled growth, improved physiological processes, and healthier wild roses.

From April to June 2023, wild roses were relocated from their natural habitats to the laboratory, considering their prevalence and suitability as *Diplolepis* species hosts. Chosen *Rosa* species encompassed *R. canina, R. rubiginosa, R. gallica*, and *R. andegavensis*. Transplanted into 20x20x25 pots with horticulture soil and sand, the roses were watered twice a week. For gall inducer oviposition, vigorous *Rosa* specimens were identified, covered with mesh tulle nets to contain the inducers. Non-galled branches (1 to 3) were marked for growth measurements post-oviposition. Similarly, control roses without of galls were chosen for comparison. Using a fine thread as reference, measurements spanned from the marked points to the branches apical shoot meristems. Length measurements were performed at least once weekly.

Our chosen gall inducers were *D. rosae* and *D. mayri*, both belonging to Cynipidae (Hymenoptera) exclusively associated with wild roses. They induce multilocular galls, each harbouring numerous chambers housing a single gall inducer larva. Females emerge in May-June and lay eggs in fresh, unopened wild rose buds. Gall growth, noticeable within weeks, extends until August-September, reaching maturity with final larval stages. Over winter, larvae pupate, emerging the subsequent spring (May-June). Structural contrast arises in the moss-like, elongated surface of *D. rosae* galls versus the shorter, spine-like nature of *D. mayri* galls.

In April and May 2023, we collected *D. rosae* and *D. mayri* galls near Cluj-Napoca, Romania. Galls were kept in plastic cups until female gall inducers emerged. Once emerged, females were put under the mesh tulle covering the rose pots. Following successful oviposition (Figure 1) and appearance of young galls, we photographed the galls weekly using a ruler as reference. The largest diameter of these galls was subsequently measured from these photos using Image J software (Schneider *et al*., 2012). Analyses were run using the R Studio (RStudio Team, 2023) with R v4.2.3 (R Core Team, 2023).

**Figure 1.**
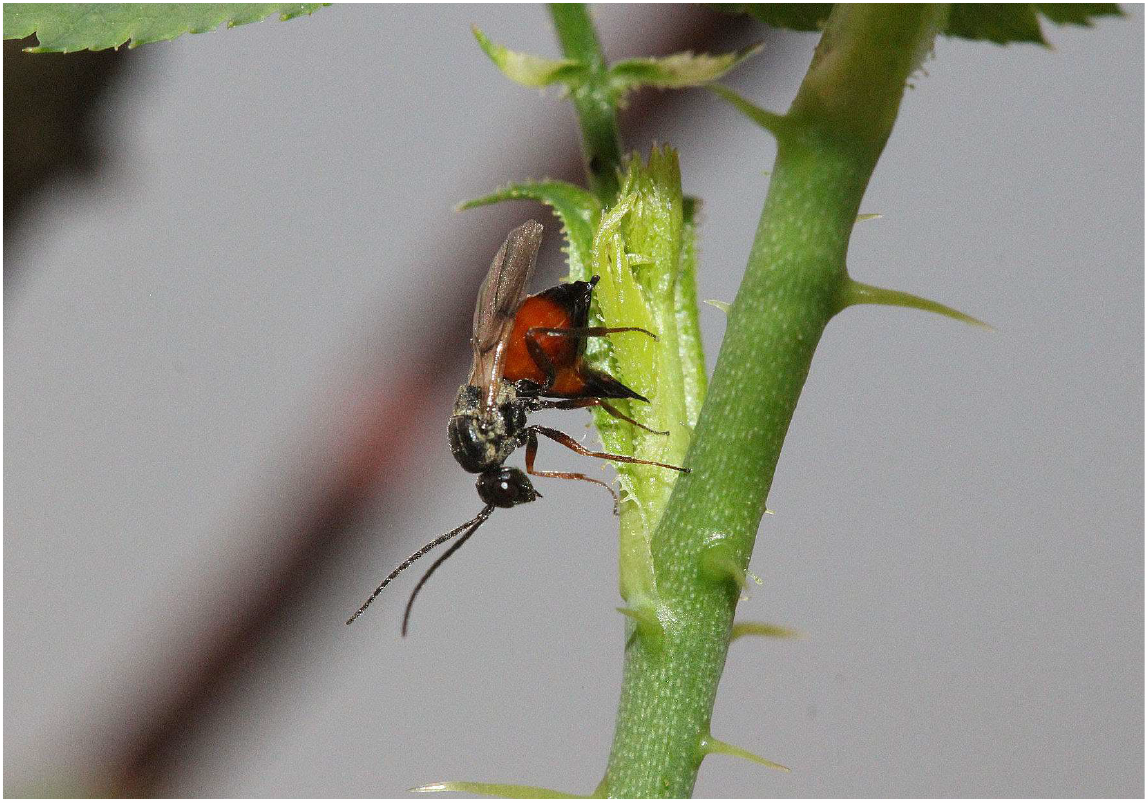
Oviposition by a *Diplolepis rosae* female on a sweet briar (*Rosa rubiginosa*).

## Results and Discussion

About 90% of transplanted roses prevailed, with only 28% successfully developing galls. Galling success was confined to *R. rubiginosa* individuals. Within the 1-hour observation windows, 26 ovipositions were noted, leading to 16 *D. mayri* and 14 *D. rosae* gall initiations. While 7 *D. mayri* and 45 *D. rosae* females were placed under the meshes, indicating greater effectiveness of *D. mayri. D. mayri* exhibited a gall-inducing efficiency of 228%, surpassing *D. rosae*’s 31%.

Although ungalled roses displayed slightly greater growth than galled ones, the discrepancy in plant growth between galled and ungalled sweet briars was not significant (linear mixed effects model: χ2=1.5, df=1, p=0.22, fixed variables: growth; random variables: bushID, date) (Figure 2A). Gall growth of both *D. mayri* (Figure 2B) and *D. rosae* (Figure 2C) was logarithmic over 42 days (between 2023.06.05 and 2023.07.17, significant estimates in a logarithmic NLS model). Notably, plant vigor impacted rose gall diameter in one case: *D. rosae* showed a significantly larger increase in diameter based on branch length and growth than *D. mayri* (ANCOVA: est.=9.75, SE=2.16, t=4.52, p= 0.001) (Figure 2 C and D).

**Figure 2.**
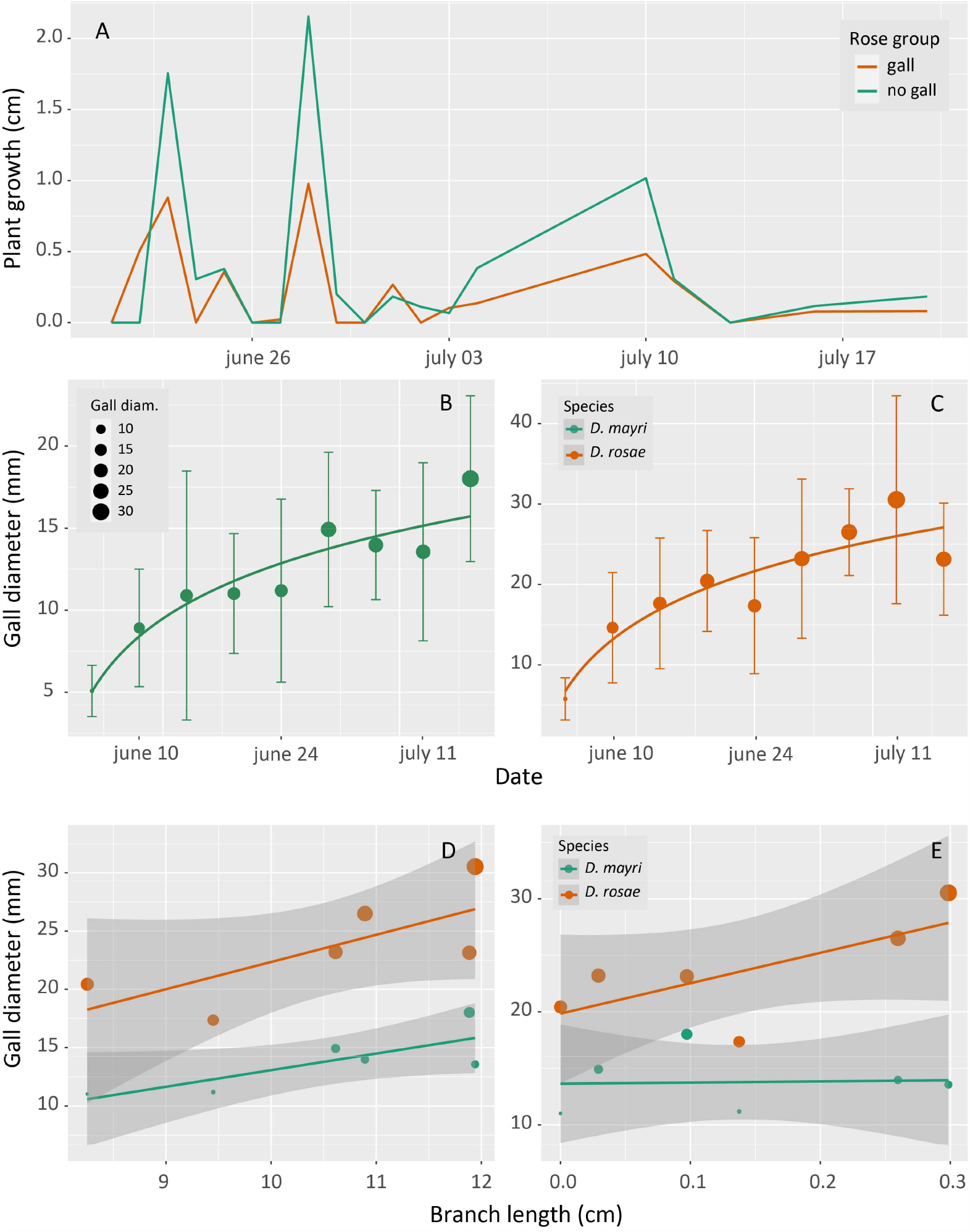
A) Plant growth (cm) of galled (N=6) and ungalled (N=6) sweet briars (*Rosa rubiginosa*). B and C) Gall growth of *Diplolepis mayri* (N=6) and *D. rosae* (N=9) over 42 days (2023.06.05 - 2023.07.17) formed on two selected sweet briars (*Rosa rubiginosa*). D and E) Plant vigour effect on the gall development over 42 days. Plant vigour expressed as the branch length or growth (cm). Gall development expressed as the diameter of the gall (mm).

We have successfully established a novel laboratory community comprising a woody plant and cynipid gall inducers. The successful induction of galls by two rose gall wasps on transplanted sweet briars is particularly noteworthy. This system enabled the study of host plant vigour’s impact on gall inducer galls. The forthcoming second laboratory generation of gall inducers and their galls will facilitate greater sample collection, enhancing the feasibility of proteomic and transcriptomic investigations. Our future plans include incorporating more wild roses and *Diplolepis* gall inducer species next year, as we’ve only explored a third of the European occurrences thus far.

## Acknowledgements

We thank Balázs Robert Zoltán for his help with the rose growth facilities and Mátis Attila for the identification of wild rose species.

## Data availability statement

The data that support the findings of this study are available on request from the corresponding author. The data are not publicly available due to privacy or ethical restrictions.

